# Performance of serological antibody tests for bovine tuberculosis in cattle from infected herds in Northern Ireland

**DOI:** 10.1101/235184

**Authors:** L. McCallan, C. Brooks, C. Couzens, F. Young, A.W. Byrne, J. McNair

## Abstract

The ability to accurately identify infected hosts is the cornerstone of effective disease control and eradication programs. In the case of bovine tuberculosis, caused by infection with the pathogen *Mycobacterium bovis*, accurately identifying infected individual animals has been challenging as all available tests exhibit less than 100% discriminatory ability. Here we assess the utility of three serological tests and assess their performance relative to skin test (Single Intradermal Comparative Cervical Tuberculin; SICCT), gamma-interferon (IFNγ) and post-mortem results in a Northern Ireland setting. Furthermore, we describe a case-study where one test was used in conjunction with statutory testing.

Serological tests using samples taken prior to SICCT disclosed low proportions of animals as test positive (mean 3% positive), despite the cohort having high proportions with positive SICCT test under standard interpretation (121/921; 13%) or IFNγ (365/922; 40%) results. Furthermore, for animals with a post-mortem record (n=286), there was a high proportion with TB visible lesions (27%) or with laboratory confirmed infection (25%). As a result, apparent sensitivities within this cohort was very low (≤15%), however the tests succeeded in achieving very high specificities (96-100%). During the case-study, 7/670 (1.04%) samples from SICCT negative animals from a large chronically infected herd were serology positive, with a further 10 animals being borderline positive (17/670; 2.54%). 9/17 of these animals were voluntarily removed, none of which were found to be infected (-lesions/-bacteriology) post-mortem; 1 serology test negative animal was subsequently lesion+ and *M bovis* confirmed at slaughter.

**Importance:** Eradication of bovine tuberculosis (bTB; caused by *Mycobacterium bovis*) has remained elusive in a number of countries despite long-term coordinated test and cull programs. This can partially be explained by the limitations of available statutory tests; therefore supplementary test platforms that identify additional infected animals would be of significant utility. Overall, during our study three serological tests did not disclose a high proportion of animals as infected in high-risk cattle herds, and exhibited limited ability to disclose animals that were positive to the statutory skin test, the gamma interferon test (IFNγ), or were post-mortem confirmed with *M. bovis*. These serological tests could be used in a supplementary fashion to the statutory tests in particular circumstances; but may be of limited advantage where parallel use of IFNγ and skin testing is performed, as these tests together tended to disclose the majority of animals with post-mortem evidence of infection in our study cohort.

## 1. Introduction

Bovine tuberculosis is a globally distributed infectious disease. The impact of infection in cattle at the national and local level can be profound (1, 2). For example, in Northern Ireland legislation is in place, supported by the United Kingdom and the European Union, to control this disease with the eventual aim of total eradication (3). In practical terms, disease control across Northern Ireland is implemented through the single intradermal comparative cervical tuberculin (SICCT) test and through carcass inspection at abattoirs where cattle are slaughtered (3). Animals identified as skin test reactors, either by standard or severe test interpretation, are removed for slaughter by compulsory order and examined post-mortem. Furthermore, all animals slaughtered at abattoirs in Northern Ireland are examined for the presence of tuberculous lesions. Clinical material collected during meat inspection is cultured for the presence of acid fast bacteria with subsequent identification of species and strain type (4).

Despite the introduction of statutory control measures to identify and remove infected cattle, bovine TB is a persistent problem in Northern Ireland (5). The epidemiology of disease is complicated by the presence of infection in wildlife (6, 7), and the potential confounding effects of concurrent infections (8, 9, 10). Current diagnostic tests applied to cattle are not sufficiently sensitive to identify all infected animals and to remove them before infection is spread (11-14). This is despite the introduction and widespread use of the interferon gamma release assay (IFNγ; 15) to augment the bovine TB testing regime and to support the front line tests (16). In combination, meat inspection, the skin test and IFNγ tests will identify a significant number of infected cattle, but not all (17). It is therefore important to investigate and validate tests or improved test strategies that will broaden the capacity to identify infected animals.

The development of serology based assays has been very useful for diagnosis where there is a Th2 type immune response. Such assays can be high throughput, relatively inexpensive and blood samples can be submitted to the laboratory a substantial time after they have been taken from the animal. However, with certain diseases a Th1 type immune responses predominates and antibody tests are largely inappropriate. This is usually the case with bovine TB when following infection, the immune response is influenced by T-cells that direct and maintain a response dominated by IFNγ release (18). Should disease progress and the burden of infection increase then the immune response changes subtly to a Th2 type where B-cells release antibody (19). In this situation and in the absence of cell mediated responses that can be exploited using the skin test or the IFNγ assay, an antibody assay may prove useful in the diagnosis of disease. In order to assess the role of antibody tests within a disease control programme that is already based on cell mediated responses, we instigated a study that was centred on bovine TB diseased cattle and at-risk herds. In the study reported here, we compared results from two blind tested antibody assays with the skin test, post-mortem examination, culture confirmation and the IFNγ assay in order to define the utility of serology as a potential diagnostic test. We also report on a case-study where one of the serological tests was used in a large herd where there was a recent chronic history of bTB, and where statutory tests were failing to clear infection.

## 2. Materials and Methods

### 2.1. Study cohort

Samples intended for analysis were taken from cattle selected from Northern Ireland herds that were deemed to have a bovine TB problem and were eligible for inclusion in the IFNγ testing scheme operated by the Department for Agriculture, Environment and Rural Affairs (DAERA), Northern Ireland (see 16, 17). Individual blood samples were taken just prior to the inoculation of tuberculins on day one of the skin test and were submitted to the laboratory within 8 hours of collection. Whole blood was removed and stimulated with antigens, to be tested later for IFNγ release. Residual whole blood was centrifuged for 15 minutes to separate plasma from blood cells. Clarified plasma samples were removed individually and stored at - 20°C.

#### The skin test and carcass inspection at abattoir

All animals included in the study were skin tested under Annex A, Council Directive 64/432/EEC using Prionics tuberculins (PPD_bovis_and PPD_avium_). Each tuberculin (0.1mL) was injected intradermally at 3000 IU (PPD_bovis_) or 2500 IU (PPD_avium_) on day one of the test. Skin thickness measurement, pre- and 72 hours post-injection was used to calculate increased skin thickness and to indicate the diagnostic outcome of the test. Skin test positive cattle were submitted for slaughter at a designated abattoir in Northern Ireland where carcass inspection was carried out to reveal the presence or absence of tuberculous lesions. Tissue samples were taken from tissues with and without tuberculous-like lesions and submitted to the culture laboratory. Information pertinent to the skin test, and abattoir inspection as well as laboratory test data was recorded onto the Animal and Public Health Information System (APHIS) operated by DAERA.

### 2.2 Laboratory procedures

#### Blinded approach to laboratory tests

Sample testing was conducted using a single blind study design in which sample information, including herd number, ear tag, other laboratory test results, was withheld from technical staff. This was achieved by assigning arbitrary codes to plasma samples upon collection. The arbitrary codes and corresponding sample information was stored in a database which was controlled by a senior technician. In compliance with data protection, information relating to herds, animals, or samples was withheld.

#### The Interferon gamma (IFNγ) test

Whole blood samples were tested for IFNγ release using the Bovigam assay (Prionics, Switzerland) accredited by the United Kingdom Accreditation Service (UKAS). The methodology has been described previously (15). Briefly, whole blood samples were received into the laboratory within eight hours of removal from the animal, stimulated overnight with Pokeweed mitogen (2ug/ml), phosphate buffered saline, PPD_bovis_, PPD_avium_ (both at 2ug/ml) and ESAT-6 (0.5ug/ml). After overnight culture at 37°C, plasma supernatant fluids were removed and stored prior to test by ELISA. The ELISA was carried out according the manufacturer’s protocol with regards to reagent dilutions, incubation times and plate wash regimes. Individual sample results were recorded if reagent control and quality assurance standards were met. Those samples with a net optical density (OD) index of 0.1 or greater were positive (net PPD_bovis_– net PPD_avium_) and those less than 0.1 OD units were negative.

#### Selection of serological tests

Tests to be evaluated were based on commercial availability and/or through fulfilling the tender to test samples via a public tender established by AFBI. Two test providers were identified (see below) who satisfied the tender requirements.

#### The IDEXX ELISA for antibodies

IDEXX *M. bovis* ELISA kits were purchased from the manufacturer and the assay was carried out according to the manufacturer’s protocol. The IDEXX ELISA is a commercially available kit. This ELISA has a 96 well microtitre plate format that detects antibodies to two *Mycobacterium tuberculosis* complex antigens (MPB70 and MPB83) known to be serological targets in *Mycobacterium bovis (M. bovis)* infections. Briefly, plasma samples were diluted to 1 in 50 in PBS and tested in duplicate. One hundred microliters of reagents were added to wells in duplicate and incubated for 60 minutes then washed 6 times. Assay positive and negative test control reagents were used to validate each microtitre plate and provided data to calculate the test result [sample / positive ratio (S/P ratio)]. Test results were interpreted as follows: an S/P ratio greater or equal to 0.30 was considered positive and a ratio less than 0.3 was negative.

#### The Enfer provisioned antibody assay

An Enfer provisioned assay was carried out by Enfer staff at their Naas laboratories (Enfer ltd, Naas, Co Kildare). All tests were blinded, with no information on the epidemiological situation (e.g. within herd prevalence) from which animals were selected provided to Enfer. It should be noted that this Enfer multiplex antibody assay is not a commercially available as a standalone kit, but testing was provided in fulfilment of commercial services as part of a commercial tender to AFBI. The basis for this assay methodology have been described previously (20). For this project, the defined antigens used in this assay were MPB83, ESAT-6, CFP-10 and MPB70. Two antigen combinations were assessed; these combinations were positive to either MPB70 with MPB83 (Enfer-A), or MPB70, MPB83, ESAT-6 and CFP10 (Enfer-B). Enfer scientific printed the bespoke multiplex according to the tender requirements, and carried out the screening, utilising bespoke software to read the multiplex plates (20). It should also be noted that the antibody test set-up did not include protein fusions and cocktails, which may be used during other Enfer tests (J. Clarke, pers. comm.). Plasma samples were diluted to 1 in 250 (in Enfer sample buffer A) and added to each well and incubated and agitated for 30 minutes. After washing, horseradish conjugated anti-bovine immunoglobulin was added, incubated and washed again. Substrate was added and signals were captured during a 45 second exposure stored as relative light units. The manufacturer recommends that a positive result is recorded when a minimum of any two antigens are test positive.

#### Laboratory confirmatory tests for Mycobacteria

Clinical samples removed from animals at slaughter were submitted to the containment level 3 laboratory for preparation, decontamination and inoculation onto solid and liquid media. Culture procedures at the Statutory TB Laboratory at the Agri-food and Biosciences Institute have been described extensively previously (e.g.17, 21). Tissue structure was disrupted using either ribolysation or grinding with sterile sand in a pestle and mortar. Prior to inoculation, clinical samples were decontaminated using 5% oxalic acid for a maximum of 30 min and washed twice with sterile PBS. Samples were then inoculated onto Lowenstein-Jensen and Stonebrink slopes, as well as into MGIT culture vessels containing PANTA. At 56 days post inoculation, cultures were examined for the presence of acid fast mycobacteria and if present were further analysed using a spoligotype method (22) to identify mycobacterial species and sub-type. Also, a selection of tissues that were lesion positive were fixed in neutral buffered formalin solution and prepared for additional histological examination.

### 2.3 Analysis

The relationship between the test status and the independent variables was modelled throughout using binary logit regression models. A random effect for herd id (to account for potential clustering effects) was included if significant and was tested using a likelihood ratio test.

Throughout we estimated the Area Under the receiver operator Curve (AUC) as an assessment of the ability of the serological test to discriminate between (apparent) infection states. The AUC is measured on a continuous scale from 0 to 1; an AUC of 0.5 is no better than random, with values >0.7 considered an “adequate” diagnostic (23). Apparent sensitivity, specificity, positive predictive value and negative predictive value was calculated and reported against non-gold standards of infection status.

Each diagnostic was compared against the skin test (SICCT) result, IFNγ test result and post-mortem status of the animal, giving apparent/relative performance indices. We also used the definition adopted by Whelan et al, (24) to define “true” infection status. In this case, infection was defined by an animal being positive to the skin test (SICCT standard interpretation), having a visible lesion at slaughter and having a bacteriological confirmation result (positive to histology and/or microbiological culture). Being free of infection, negative animals were negative to SICCT, without lesions at slaughter and without post-mortem bacteriological confirmation. In addition, we used a combination of IFNγ, SICCT, VL and confirmation, to assess the relative performance of the serology tests.

We used binary logit models to assess whether there was any association between animal sex, age at blood test sample, breed type and the probability of a positive serological test results being disclosed. Age was modelled both as a raw continuous variable and as a log-transformed predictor.

Throughout, the dataset was organised using Microsoft excel, while all statistical analysis was undertaken using Stata version 14 (Stata Corp., Texas, USA, 2015).

### 2.4 A problem herd based case study

A case study centred on a relatively large (approximately 1000 cattle over the period) dairy herd was carried out to assess the utility of antibody detection where animals were known to be infected and resolution of the problem was proving to be difficult. This particular herd had a seemingly intractable chronic bovine TB problem which originated between 2002 and 2004. Initially, a relatively small number of bovine TB breakdowns were recorded with subsequent confirmation of infection caused by *Mycobacterium bovis*. From 2008 onward, the rate of skin test positive cattle increased significantly with up to 148 skin test positive animals identified as well as 2 cases of lesions at routine slaughter, i.e. skin test negative cattle sent for slaughter with confirmed tuberculous lesions disclosed during carcass inspection. Given the disease history of this herd following routine TB diagnostic investigations, high risk cohorts of cattle within this herd were blood sampled and tested for the presence of antibodies to *M. bovis* using IDEXX serology (OIE approved) in 2016. The fundamental rationale was that detecting antibody in cattle that were skin test negative may indicate the presence of infection in animals that were considered to be anergic, that is, unresponsive to cell mediated tests such as the skin test and IFNγ assay.

## 3. Results

### Agreement and comparison

Overall, there were 922 animals with test result data; all animals had test results for IFNγ and IDEXX, 921 had SICCT, Enfer-A and Enfer-B results, while 286 animals had a post-mortem result. These animals came from 64 herds with recent bTB breakdowns, with a mean of 14.39 animals sampled per herd (Median: 9.5; Std. Dev.: 13.39; Range: 1-76).

There was significant (p<0.001) moderate agreement between the three tests ranging from a kappa of 0.40 (IDEXX and Enfer-B) to 0.55 (Enfer-A and Enfer-B). Of the animals with visible lesions found at post-mortem, the proportions deemed positive were not significantly different between the serological test types (McNemar’s test: Enfer-A vs. IDEXX: p=0.65; Enfer-B vs. IDEXX: p=0.16; Enfer-B vs Enfer-A: p=0.18). Similarly, there were no difference between test types, when using bacteriological confirmation as the infection status diagnostic (p>0.25).

### Serology test performance in comparison with single or combined diagnostic techniques

The relative performance of the serological tests in comparison with single ante-mortem diagnostics (Table1), post-mortem diagnostics (Table 2) and combined tests (Table 3 and Table 4) are presented below. Relative to single ante-mortem tests (mean test prevalence 27%; Table 1), the serological tests did not disclose a high proportion of test-positive animals (mean 3% positive). This resulted in the tests exhibiting low apparent sensitivities, averaging 5.73% (range: 4.13% - 9.09%). However, the apparent specificities were always very high, with a mean of 97.82% (96.40% - 99.50%). While there was a significant positive relationship between serological test result and statutory ante-mortem outcome, the discriminatory ability of the tests were always poor (mean AUC: 0.52).

**Table 1:**
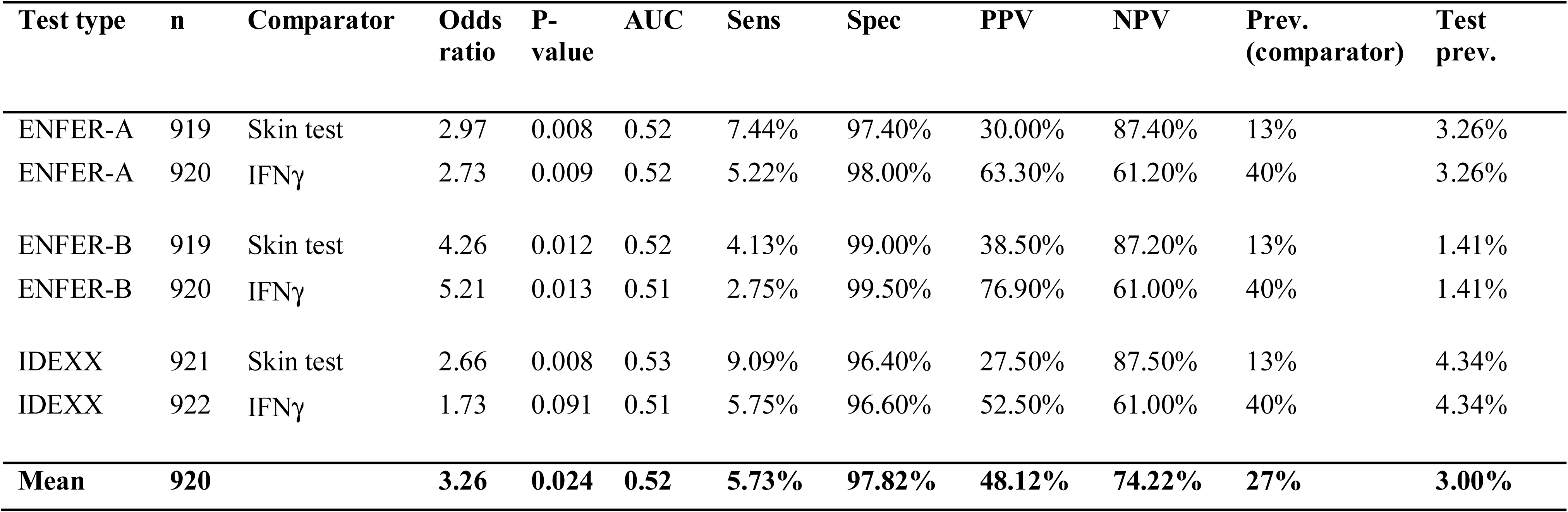
The relative performance of serological tests against statutory ante-mortem tests. Skin test = SICCT standard interpretation.

**Table 2:**
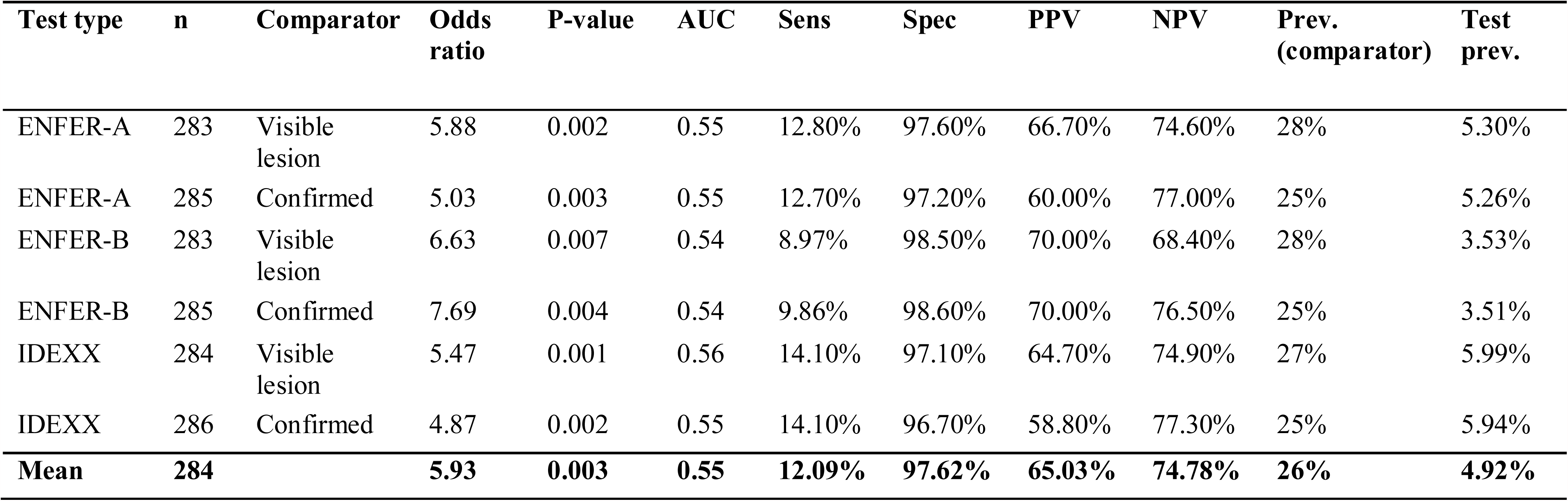
The relative performance of serological tests against statutory post-mortem diagnostic techniques.

**Table 3:** The relative performance of serological tests against a combination of statutory ante-mortem and post-mortem diagnostic techniques. Positive status animals were positive to SICCT, had a visible lesion (VL) at slaughter and had bacteriologically confirmed infection; negative status animals were negative to SICCT, VL and were not confirmed at slaughter.

| Test type | n | Comparator | Odds ratio | P-value | AUC | Sens | Spec | PPV | NPV | Prev. (comparator) | Test prev. |
| --- | --- | --- | --- | --- | --- | --- | --- | --- | --- | --- | --- |
| ENFER-A | 187 | SICCT + VL<br>+<br>CONFIRM | 14.20 | 0.019 | 0.54 | 9.09% | 99.30% | 80.00% | 78.02% | 24% | 2.67% |
| ENFER-B | 187 | SICCT + VL<br>+<br>CONFIRM | 10.39 | 0.045 | 0.53 | 6.82% | 99.30% | 75.00% | 77.60% | 24% | 2.14% |
| IDEXX | 188 | SICCT + VL<br>+<br>CONFIRM | 7.42 | 0.006 | 0.56 | 13.64% | 97.92% | 66.67% | 78.77% | 31% | 6.25% |
| **Mean** | **187** |  | **10.67** | **0.023** | **0.54** | **9.85%** | **98.84%** | **73.89%** | **78.13%** | **26%** | **3.69%** |

**Table 4:** The relative performance of serological tests against a combination of statutory ante-mortem and post-mortem diagnostic techniques. Positive status animals were positive to SICCT, Interferon-G, had a visible lesion (VL) at slaughter and had bacteriologically confirmed infection; negative status animals were negative to SICCT, IFNγ, VL and were not confirmed at slaughter.

| Test type | n | Comparator | Odds ratio | P-value | AUC* | Sens | Spec | PPV | NPV | Prev. (comparator) | Test prev. |
| --- | --- | --- | --- | --- | --- | --- | --- | --- | --- | --- | --- |
| ENFER-A | 68 | SICCT + IFNγ + VL + CONFIRM | NA | NA | 0.55 | 10.00% | 100.00% | 100.00% | 43.75% | 59% | 5.88% |
| ENFER-B | 68 | SICCT + IFNγ + VL + CONFIRM | NA | NA | 0.54 | 7.50% | 100.00% | 100.00% | 43.08% | 59% | 4.41% |
| IDEXX | 68 | SICCT + IFNγ + VL + CONFIRM | NA | NA | 0.58 | 15.00% | 100.00% | 100.00% | 45.16% | 59% | 8.82% |
| **Mean** | **68** |  |  |  | **0.55** | **10.83%** | **100.00%** | **100.00%** | **44.00%** | **59%** | **6.37%** |

Similar results were found when post-mortem diagnostic techniques were used as the apparent infection status (Table 2). Due to the low sensitivity of the serological antibody tests, the mean test prevalence was always low (mean test prevalence 4.92%) relative to the proportion of animals with lesions or post-mortem confirmed infection (mean prevalence 26%).

Using similar criteria to Whelan et al. (24) to define animals as “truly” infected and non-infected, we found that the serological tests exhibited poor sensitivity (9.09% - 13.64%; Table 3). Utilising IFNγ test results as an additional criterion (Table 4), suggested again that the serological tests exhibited low sensitivities, however the three tests achieved 100% apparent specificities.

Table 5 gives the breakdown of animal ante-mortem test results in relation of each serological test result. Overall, 8 (8/505;1.56%), 2 (2/513; 0.39%), and 17 (17/514; 3.31%) animals were ante-mortem test negative, that were deemed serologically test positive to Enfer-A, Enfer-B and IDEXX respectively.

**Table 5:**
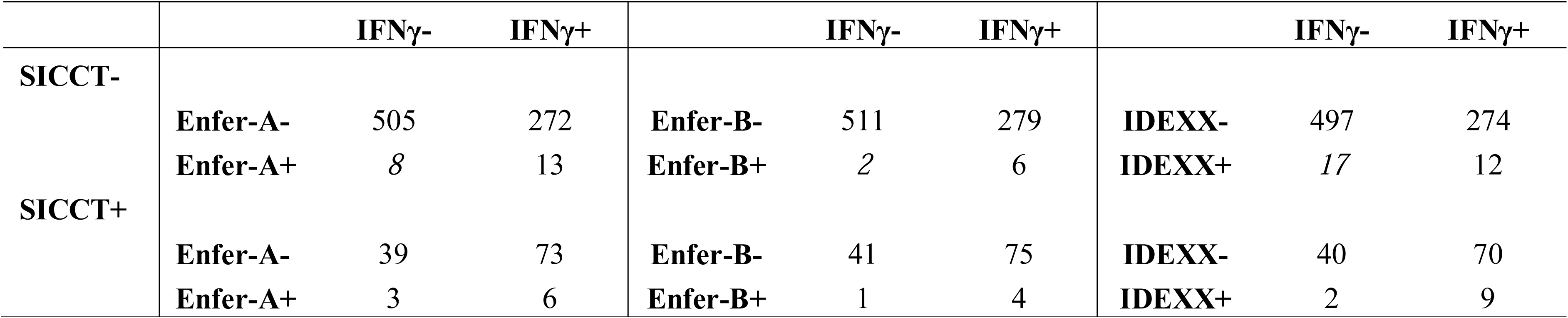
Tabulation of the relationship between serological test result, gamma interferon (IFNγ) status and skin test status. Numbers italicised represent ante-mortem negative animals that were serologically test positive.

Table 6 gives a breakdown of animals with post-mortem confirmed *M. bovis* infection, that were skin-test, IFNγ, or either skin-test/ IFNγ negative. ENFER-A and IDEXX both disclosed as positive 3/19 (15.79%) SICCT false-negative animals. The ENFER-B test disclosed two animals of these 19 animals as positive. However, none of the 14 post-mortem confirmed animals that were as IFNγ negative were found to be serologically positive. Overall, 6 of the animals with confirmed infection were missed by both SICCT and IFNγ tests (6/286; 2.10%), and none of these were disclosed using any of the serological antibody tests.

**Table 6:** Proportion of confirmed infected animals with positive serological test results, which were missed by SICCT, IFNγ, or both ante mortem bovine TB tests.

| **Confirmed infection** | **ENFER-**<br>**A** | **ENFER-**<br>**B** | **IDEXX** |
| --- | --- | --- | --- |
| **SICCT- (n)** | 3/19 | 2/19 | 3/19 |
| **(% serology positive)** | 15.79% | 10.53% | 15.79% |
| **IFNγ -** | 0/14 | 0/14 | 0/14 |
| **(% serology positive)** | 0% | 0% | 0% |
| **SICCT or IFNγ -** | 0/6 | 0/6 | 0/6 |
| **(% serology positive)** | 0% | 0% | 0% |

### Sex, age and breed associations with serological test results

There was no evidence of a significant effect of sex on the probability of an animal disclosing as serological positive across all tests (p>0.1). Similarly, there was no evidence of an age effect on the probability of animals disclosing with serological positive test (p>0.08). There was no relationship between the breed and either Enfer-A (p=0.617) or IDEXX (p=0.457) positivity. However, there was significant variation in the probability of a positive disclosure with Enfer-B (n= 683; p=0.005; Figure 1). This model included five breed types with enough samples and variation to allow the model to fit (n= 26 to 378 per breed type; Figure 1). Friesian breed cattle exhibited significantly lower odds of disclosing as an Enfer-B test positive relative to Charolais (p=0.025), Hereford (p=0.006) and Limousine (p=0.012) animals, respectively, however the difference was not significant relative to Aberdeen Angus cattle (p=0.116; see supplementary material for figure of linear predictions with associated 95%CI).

**Figure 1:**
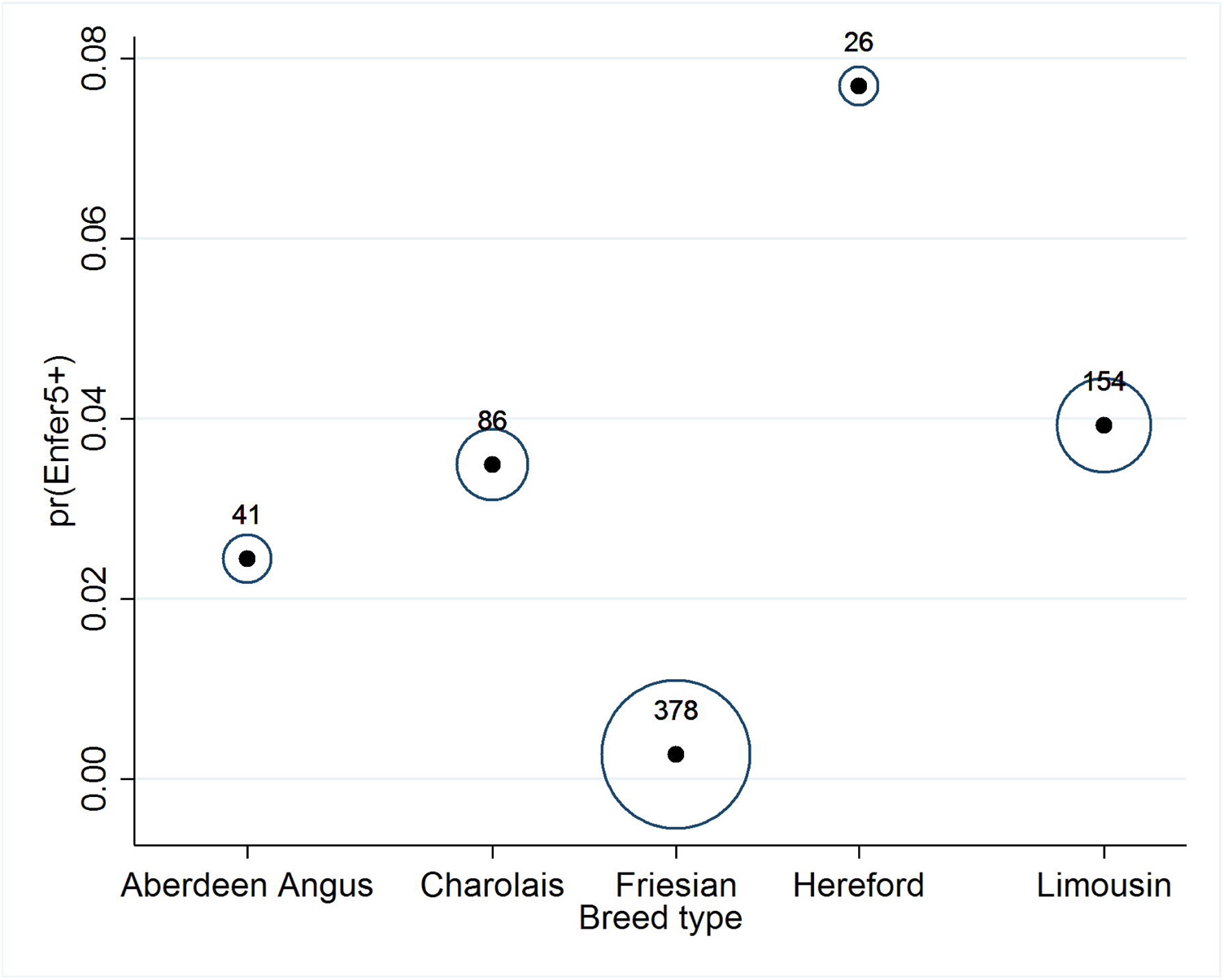
The predicted probability of an animal testing positive to the Enfer-B serological test depending on breed type. The size of the circles represents the weighted sample size for each breed type modelled; numbers represent the number of observations for each breed type.

#### A problem herd based case study

In total, 670 samples from cattle were blood sampled having been selected on the basis of being high risk cohorts of animal where the infection was most prevalent (ante-mortem negative in contact animals). Using the manufacturer’s recommended S/P ratio cut-off value of 0.3, seven samples were positive (≥ 0.3) and 663 samples were negative (≤0.3). Five samples were clearly positive (> 0.3), two samples were just above the threshold (0.340 and 0. 331) and all the remaining samples were negative. However, 17 samples had S/P ratios just below the cut-off value, ranging between 0.270 to 0.100 (Table 7).

**Table 7.** Distribution of S/P ratios for 670 samples tested for antibodies using the IDEXX ELISA for *M. bovis* in one chronically infected case-study farm in Northern Ireland.

| S/P $\geq 0.3$ | S/P < 0.3 to 0.1 | < 0.1 to 0.0 | < 0.0 to -0.094 |
| --- | --- | --- | --- |
| Positive | Negative | Negative | Negative |
| 7 | 17 | 55 | 591 |

Following release of the serology results and discussions with the herd keeper, nine animals were voluntarily surrendered for slaughter. At post-mortem examination, all cattle were designated non-visibly lesioned and clinical samples from the lung associated lymph nodes were submitted for laboratory tests. All samples were culture negative for *M. bovis.* Subsequent to this serology test based investigation, one animal which was serology negative and submitted for voluntary slaughter, was examined and found to be visibly lesioned. Subsequently, clinical samples were culture positive with *M. bovis* confirmed by spoligotype.

## 4. Discussion

During the present study, we tested three serological tests (2 versions of the ENFER multiplex, and the IDEXX AB test) for their relative performance in at-risk herds in Northern Ireland. Overall, our results suggested that the tests can achieve very high levels of apparent specificity. However, our results suggested that these tests failed to identify many animals with pathology or confirmed *M bovis* infection post-mortem. These results appear at odds with some previous studies in other populations (24, 26), with serological antibody tests being suggested as a potential useful diagnostic in certain situations (27-30). However, there has been reported lower sensitivity estimates elsewhere (e.g. 27, 31).

In comparison with previous work by our group (14), serology samples were taken prior to the SICCT tuberculin test. In work from Spain, when serology tests were evaluated prior to the tuberculin test, their performance was reduced relative to tests undertaken with samples after the tuberculin test (27). Samples taken from a cohort of animals prior to skin testing suggested that the serology tests examined exhibited a sensitivity of 23.9%-32.6% (M *bovis* Ab Test (IDEXX) & Enferplex TB assay, respectively; 27). For animals sampled post-skin test, the beneficial anamnestic effect was most pronounced 15 days post-intradermal testing, achieving sensitivity estimates of 66.7%-85.2% (27). The effect was apparent by the number of animals disclosed as serology test positive when tested prior to skin testing (10.7%; 6/56), 72hrs after skin testing (7.1%; 4/56) and 15 days after testing (57.1%; 32/56). In the current study, a small proportion of animals were disclosed as serology positive (mean 3% positive). However, during another study in Northern Ireland, we found a higher proportion of animals were disclosed as positive when prevalence was higher (86% SICCT test reactors) and testing occurred after skin testing (14). The proportion serology positive in that cohort was 39.02-62.20% positive, with apparent sensitivities relative to post-mortem confirmed infection estimated to be 68-82%. These results suggest that maximising the beneficial effects of serology testing may occur if samples are taken after skin testing. Such boosting/priming effects have been described before in cattle in a number of studies (27, 28, 32-35) and in other species also (see 36). Two antigens used in the tests assessed during the present study are known to be boosted by skin testing (MPB83 and MPB70; 35). Such effects have led to some authorities to require follow-up serology testing during statutory tests, for example with camelids in Wales (36). However, this effect reduces the utility of such a test in cattle where proactive eradication programs are in place, as skin test positive animals in Northern Ireland are routinely culled under legislation (3). Furthermore, in problem herds judicious use of severe interpretation of the SICCT (this increases sensitivity, at the cost of specificity (13)) is employed, and also INFγ tests (16; and see below), where positive animals are predominately culled to clear infection.

In the present study, a small proportion of infected but SICCT negative animals that were identified by the serological tests (2-3/19 animals; 10.53%-15.79%). This suggests that, in the absence of other ancillary testing, serological tests could be useful to identify part of this subpopulation (Whelan et al. (24)). Previous research found that of 60 truly infected SICCT negative or inconclusive animals, 53 (88.3%) were disclosed as positive using a multiplex ELISA test (24). It is hard to account for the relatively poorer detection rate in our study relative to Whelan et al. (24), but the discrepancy can partly be explained by the relatively small number of SICCT negative, *M. bovis* confirmed animals available in the present study.

Employing exact confidence intervals around the proportion, suggests significant uncertainty in our estimate (exact CI: 3.38% - 39.58%). In Northern Ireland, IFNγ is routinely used in herds with problems clearing infection (e.g. see 16, 17). We found in this study, that when IFNγ was used instead of, or in parallel with, SICCT, there were no *M. bovis* confirmed animals identified by the serological tests employed. This suggests, where both SICCT and INFy are used together, there may be limited opportunities to detect additional missed infected animals using serological tests. Casal et al. (29), however, suggests that in very high prevalence regions there may be value in parallel interpretation of cellular and antibody detection techniques to maximise sensitivity.

During the case study presented, we found that few animals were disclosed as serologically positive from a large herd with a substantial chronic bTB problem. Even with liberal interpretation of one of the serology test (IDEXX) data, few animals were removed, and all those culled were found to have no visible lesions nor could *M. bovis* be isolated from samples taken from these animals. One animal that was serologically tested, and found negative, was subsequently found to have visible lesions and confirmed for *M. bovis* at postmortem. This field application of the test in a particular problematic herd appears to corroborate our findings from the prospective study results. However, other case-studies have highlighted benefits of serology as ancillary tests in eradicating TB. For example, a red deer herd in England with a TB outbreak was cleared of infection with the use of both tuberculin testing and serological testing over a 2 year period (30). The authors suggest that without the additional removal of serologically test positive, the time to eradication may have been significantly increased as well as contributing to maintenance and potential transmission to local wildlife. O’Brien et al. (28) also describes a case-study in a goat herd where skin tests failed to identify all infected animals, with 6/20 slaughtered animals having visible lesions and positive to six *M. bovis* antigens.

Serological tests could be strategically useful in the case of anergic animals, where advanced and generalised infection is present leading to failure to respond to SICCT due to an impaired Cell mediated immunity (CMI) response (12). However, currently there is limited data on the proportion of animals that could be deemed anergic in Northern Ireland farms. Potentially, the repeated application of SICCT testing over an animal’s lifetime could lead to desensitisation (12, 37), again resulting in false negatives. When we looked at the impact of age on the probability of disclosure, we found no significant variation in our cohort. However, we did find some evidence for variation in disclosure depending on breed-type, with Friesian cattle exhibiting significantly lower probability of disclosing serology positive on one of the tests. Further research is required to ascertain whether this is a robust finding – there is significant uncertainty with the current study given the very small numbers of animals serologically test-positive. However, previous research has suggested that there may be significant variation in *M. bovis* susceptibility and pathology across breeds (38, 39), which could be partially attributed to immunological or genetic variation (40).

One potential reason for the differing outcomes from this study and some other studies using the Enfer test platform, is that there was a limited set of antigens used across the two test types (Enfer-a and Enfer-b), namely MPB70, MPB83, ESAT-6 and CFP10. The Enfer multiplex can detect antibody activity to 25 antigens in a single well in a 96-well plate array format (20). However, to make cross-comparisons, only the most commonly used antigens were used during the present study. However, such issues do not arise with the IDEXX-ab test, as it is a standard commercial kit.

## Conclusions

We have shown that three available serological tests, when applied to cattle populations with moderate prevalence and with samples taken prior to tuberculin testing, can exhibit limited apparent sensitivities but very high specificities. Serological tests can disclose additional test-positive animals when used in parallel with the skin tuberculin test. However, we found in this study, that when IFNγ was used instead of, or in parallel with, SICCT, there were no *M. bovis* confirmed animals identified by the serological tests employed. This suggests, where both SICCT and IFNγ are used together, there may be limited opportunities to detect additional missed infected animals via the serological tests examined when samples were taken prior to skin testing. From a perspective of a country with an ongoing extensive eradication scheme, future strategic use of serology may be limited to: 1. extreme cases of very large breakdowns within herds leading to high within herd bTB prevalence, 2. in problem herds where IFNγ testing is unavailable, and 3. chronically infected herds where blood samples are taken after tuberculin testing to maximise sensitivity gained from any anamnestic effects.

## Data availability

All data was provided through the APHIS dataset, for which the data controller is DAERA. All data from which inferences were made are provided within the paper, raw test data has been deposited in an online repository (25). Additional information on these data is available from DAERA, Northern Ireland (https://www.daera-ni.gov.uk/access-information-0;daera.informationmanager@daera-ni.gov.uk) and would be subject to appropriate Data Protection regulations (UK) in relation to individual farmers/herds.

## Acknowledgements

This study was funded by the Department of Agriculture, Environment and Rural Affairs (DAERA) as part of the Evidence and Innovation Strategy under the grant “An assessment of commercially available serological tests for the detection of cattle infected with bovine tuberculosis” (grant no.: 15/3/09; Project Leaders: Dr. J. McNair and Dr. A. Byrne).

## References

Zinsstag J, Schelling E, Roth F, Kazwala RR. 2006. Economics of bovine tuberculosis. *In* Thoen CO (Ed.), Mycobacterium bovis infection in animals and humans. 2nd edition. Blackwell Publishing, pp. 68–83

O’Reilly LM, Daborn CJ 1995. The epidemiology of *Mycobacterium bovis* infections in animals and man: a review. Tuber Lung Dis 76:1–46.

Abernethy DA, Denny GO, Menzies FD, McGuckian P, Honhold N, Roberts AR. 2006. The Northern Ireland programme for the control and eradication of *Mycobacterium bovis*. Vet Microbiol 112:231–237.

Skuce RA, McDowell SW, Mallon TR, Luke B, Breadon EL, Lagan PL, McCormick CM, McBride SH, Pollock JM. 2005. Discrimination of isolates of *Mycobacterium bovis* in Northern Ireland on the basis of variable numbers of tandem repeats (VNTRs). Vet Rec 157:501.

Abernethy DA, Upton P, Higgins IM, McGrath G, Goodchild AV, Rolfe SJ, Broughan JM, Downs SH, Clifton-Hadley R, Menzies FD, De la Rua-Domenech R. 2013. Bovine tuberculosis trends in the UK and the Republic of Ireland, 1995-2010. Vet Rec 172:312–312.

Palmer MV. 2007. Tuberculosis: a reemerging disease at the interface of domestic animals and wildlife. In Wildlife and Emerging Zoonotic Diseases: The Biology, Circumstances and Consequences of Cross-Species Transmission (pp. 195-215). Springer Berlin Heidelberg.

Byrne AW, White PW, McGrath G, O’Keeffe J, Martin SW. 2014. Risk of tuberculosis cattle herd breakdowns in Ireland: effects of badger culling effort, density and historic large-scale interventions. Vet Res 45:1.

Claridge J, Diggle P, McCann CM, Mulcahy G, Flynn R, McNair J, Strain S, Welsh M, Baylis M, Williams DJ. 2012. Fasciola hepatica is associated with the failure to detect bovine tuberculosis in dairy cattle. Nature Comm 3:853.

Byrne AW, Guelbenzu-Gonzalo M, Strain SAJ, McBride S, Graham J, Lahuerta-Marin A, Harwood R, Graham DA, McDowell S. 2017. Assessment of concurrent infection with bovine viral diarrhoea virus (BVDV) and *Mycobacterium bovis:* a herd-level risk factor analysis from Northern Ireland. Preventive Vet Medicine.

Álvarez J, De Juan L, Bezos J, Romero B, Sáez JL, Marqués S, Domínguez C, Mínguez O, Fernández-Mardomingo B, Mateos A, Domínguez L. 2009. Effect of paratuberculosis on the diagnosis of bovine tuberculosis in a cattle herd with a mixed infection using interferon-gamma detection assay. Vet Microbiol 135:389–393.

Pollock JM, Welsh MD, McNair J (2005) Immune responses in bovine tuberculosis: towards new strategies for the diagnosis and control of disease. Vet Immunol Immunopathol 108:37–43

de la Rua-Domenech R, Goodchild AT, Vordermeier HM, Hewinson RG, Christiansen KH, Clifton-Hadley RS. 2006. Ante mortem diagnosis of tuberculosis in cattle: a review of the tuberculin tests, γ-interferon assay and other ancillary diagnostic techniques. Res Vet Sci 81:190–210.

Lahuerta-Marin A, Milne G, McNair J, Skuce R., McBride S., Menzies F., McDowell S.J.W., Byrne A.W., Handel I.G., Bronsvoort M.B.C. 2018. Bayesian Latent Class estimation of sensitivity and specificity parameters of diagnostic tests for bovine tuberculosis in chronically infected herds in Northern IrelandBayesian Latent Class estimation of Sensitivity and Specificity parameters of diagnostic tests for bovine tuberculosis in chronic herds-Northern Ireland. Vet J in press.

McCallan L, Brooks C, Couzens C, Young F, McNair J, Byrne AW. 2017. Assessment of serological tests for diagnosis of bovine tuberculosis. Vet Rec 181: 90

Welsh MD, McNair J, McDowell SWJ, Buchanan J, Hill R, McBride SH, Clarke N. 2008. The Northern Ireland Interferon-gamma (IFN-g) Testing Programme. Cattle Practice 16:136–139

Lahuerta-Marin A, Gallagher M, McBride S, Skuce R, Menzies F, McNair J, McDowell SW, Byrne AW. 2015. Should they stay, or should they go? Relative future risk of bovine tuberculosis for interferon-gamma test-positive cattle left on farms. Vet Res 46:1.

Lahuerta-Marin A, McNair J, Skuce R, McBride S, Allen M, Strain SA, Menzies FD, McDowell SJ, Byrne AW. 2016. Risk factors for failure to detect bovine tuberculosis in cattle from infected herds across Northern Ireland (2004-2010). Res Vet Sci 107:233–239.

Welsh MD, Cunningham RT, Corbett DM, Girvin RM, McNair J, Skuce RA, Bryson DG, Pollock JM. 2005. Influence of pathological progression on the balance between cellular and humoral immune responses in bovine tuberculosis. Immunol 114:101–111.

Pollock JM, Neill SD. 2002. *Mycobacterium bovis* infection and tuberculosis in cattle. Vet J 163:115–127.

Whelan C, Shuralev E, O’Keeffe G, Hyland P, Kwok HF, Snoddy P, O’Brien A, Connolly M, Quinn P, Groll M, Watterson T. 2008. Multiplex immunoassay for serological diagnosis of *Mycobacterium bovis* infection in cattle. Clin Vac Immunol 15:1834–1838.

Stewart LD, McNair J, McCallan L, Gordon A, Grant IR. 2013. Improved detection of *Mycobacterium bovis* infection in bovine lymph node tissue using immunomagnetic separation (IMS)-based methods. PloS one 8:e58374.

Kamerbeek J, Schouls LE, Kolk A, Van Agterveld M, Van Soolingen D, Kuijper S, Bunschoten A, Molhuizen H, Shaw R, Goyal M, Van Embden J. 1997. Simultaneous detection and strain differentiation of Mycobacterium tuberculosis for diagnosis and epidemiology. J Clin Microbiol 35:907–14.

Hosmer DW, Lemeshow S. 2004. Applied logistic regression. John Wiley & Sons.

Whelan C, Whelan AO, Shuralev E, Kwok HF, Hewinson G, Clarke J, Vordermeier HM. 2010. Performance of the Enferplex TB assay with cattle in Great Britain and assessment of its suitability as a test to distinguish infected and vaccinated animals. ClinVaccine Immunol 17:813–817.

McCallan L, Brooks C, Couzens C, Young F, McNair J, Byrne AW. 2017. Data from Performance of serological antibody tests for bovine tuberculosis in cattle from infected herds in Northern Ireland. Figshare. https://figshare.com/s/db7f5956d1094066180a DOI: 10.6084/m9.figshare.5705893

Waters WR, Buddle BM, Vordermeier HM, Gormley E, Palmer MV, Thacker TC, Bannantine JP, Stabel JR, Linscott R, Martel E, Milian F. 2011. Development and evaluation of an enzyme-linked immunosorbent assay for use in the detection of bovine tuberculosis in cattle. ClinVaccine Immunol 18:1882–1888.

Casal C, Infantes JA, Risalde MA, Díez-Guerrier A, Domínguez M, Moreno I, Romero B, de Juan L, Sáez JL, Juste R, Gortázar C. 2017. Antibody detection tests improve the sensitivity of tuberculosis diagnosis in cattle. Res Vet Sci 112:214–21.

O’Brien A, Whelan C, Clarke JB, Hayton A, Watt NJ, Harkiss GD. 2017. Serological Analysis of Tuberculosis in Goats by Use of the Enferplex Caprine TB Multiplex Test. Clin Vaccine Immunol 24:e00518-16.

Busch F, Bannerman F, Liggett S, Griffin F, Clarke J, Lyashchenko KP, Rhodes S. 2017. Control of bovine tuberculosis in a farmed red deer herd in England. Vet Rec 180:68.

Wood PR, Corner LA, Rothel JS, Ripper JL, Fifis T, McCormick BS, Francis B, Melville L, Small K, De Witte K, Tolson J. 1992. A field evaluation of serological and cellular diagnostic tests for bovine tuberculosis. Vet Microbial 31:71–79.

Waters WR, Palmer MV, Thacker TC, Bannantine JP, Vordermeier HM, Hewinson RG, Greenwald R, Esfandiari J, McNair J, Pollock JM, Andersen P. 2006. Early antibody responses to experimental *Mycobacterium bovis* infection of cattle. Clin Vaccine Immunol 13:648–54.

Waters WR, Maggioli MF, McGill J, Lyashchenko KP, Palmer MV. 2014. Relevance of bovine tuberculosis research to the understanding of human disease: Historical perspectives, approaches and immunologic mechanisms. Vet Immunol Immunopathol 159:113–132.

Waters WR, Thacker TC, Nelson JT, DiCarlo DM, Maggioli MF, Greenwald R, Esfandiari J, Lyashchenko KP, Palmer MV. 2014. Virulence of two strains of *Mycobacterium bovis* in cattle following aerosol infection. J Com Pathol 151:410–419.

Waters WR, Palmer MV, Stafne MR, Bass KE, Maggioli MF, Thacker TC, Linscott R, Lawrence JC, Nelson JT, Esfandiari J, Greenwald R. 2015. Effects of serial skin testing with purified protein derivative on the level and quality of antibodies to complex and defined antigens in *Mycobacterium* bovis-infected cattle. Clin Vaccine Immunol 6:641–649.

de la Rua-Domenech R, Rhodes S, Rolfe S, Vordermeier M. 2017. The anamnestic boost effect of the skin test on antibody responses to Mycobacterium bovis in camelids - summary of evidence. http://www.alpacatb.com/anamnestic%20antibody%20response%20(scientific%20evidence)%20(2).pdf

Monaghan ML, Doherty ML, Collins JD, Kazda JF, Quinn PJ. 1994. The tuberculin test. Vet Microbiol 40:111–24.

Ameni G, Aseffa A, Engers H, Young D, Gordon S, Hewinson G, Vordermeier M. 2007. High prevalence and increased severity of pathology of bovine tuberculosis in Holsteins compared to zebu breeds under field cattle husbandry in central Ethiopia. Clin Vaccine Immunol 14:1356–1361.

Wright DM, Allen AR, Mallon TR, McDowell SW, Bishop SC, Glass EJ, Bermingham ML, Woolliams JA, Skuce RA. 2013. Field-isolated genotypes of *Mycobacterium bovis* vary in virulence and influence case pathology but do not affect outbreak size. PloS one 8:e74503

Allen AR, Minozzi G, Glass EJ, Skuce RA, McDowell SWJ, Woolliams JA, Bishop SC. 2010. Bovine tuberculosis: the genetic basis of host susceptibility. P R Soc Lond B: Biol Sci rspb20100830.

